# The time course of updating in running span

**DOI:** 10.1101/522391

**Authors:** Shraddha Kaur, Dennis Norris, Susan E Gathercole

**Affiliations:** Medical Research Council Cognition and Brain Sciences Unit, University of Cambridge

**Keywords:** running span, working memory updating, cognitive demand, working memory

## Abstract

Running span can be performed by either passively listening to to-be-remembered items or actively updating the target set during presentation. This choice of strategy is influenced by the rate of presentation in the task. Previous research suggests that the active updating process is demanding and time-consuming. It is favored at relatively slow rates of presentation, while the passive strategy is more successful when applied at fast rates. In two experiments the time course of resource demand during task performance and its sensitivity to presentation rate was examined. We hypothesized that running span imposes a high cognitive load only when active updating is employed. Participants performed running span simultaneously with a spatial reaction time (RT) task, and RTs on the concurrent task were used to index the resource demands of the memory task. A slow-paced running span exhibited a large overall resource demand in comparison with the serial recall tasks (Experiment 1) and fast-paced running span (Experiment 2). This demand was observed from the position in the list from which participants are presumed to start updating, suggesting a cognitive shift to a demanding mode of updating. In addition, a demand burst was found approximately 1000ms following item onset at these later positions. These data establish that the process of active updating in running span task is slow and cognitively demanding and indicate that this limits its application during fast presentation rates.

---

Running span is a complex working memory task that places a unique demand on the maintenance of serially ordered information. Only the latest information is relevant for recall at any given point in the task, but the timing of this query is unpredictable. Participants appear to keep track of this relevant information by actively updating the target recall set in working memory (WM) when new information is presented (Bunting, Cowan & Saults, 2006; Hockey, 1973; Morris and Jones, 1990). Specific cognitive processes supporting updating have been investigated in tasks that involve updating of individual items on the basis of item characteristics such as the most recent exemplar from particular semantic categories (Oberauer, 2018; Lewis-Peacock, Kessler & Oberauer, 2018; Kessler & Oberauer, 2014; 2015). It is unclear whether the same operations can be applied to running span, in which participants attempt to recall the *n* most recent items in a lengthy sequence. The aim of the present study was to provide the first fine-grained temporal analysis of the cognitive demands of the updating process in running span. We did this by using performance on a concurrent reaction time task to index resource consumption during the process of updating.

In a typical running span experiment participants are presented a sequence of variable length and are asked to recall the last *n* items of the sequence in serial order. In the first substantial experimental examination of running span, Hockey (1973) identified two strategies that participants could use to perform the task: passive listening and active processing. The passive mode involves receiving incoming items without engaging in any additional processing or actively attempting to update the recall set. Cowan and colleagues proposed that incoming information is stored as a sensory trace in the first instance and then converted to a categorical form appropriate for recall at the end of the list (Bunting et al., 2006; Cowan et al., 2005). When spoken lists are presented, retrieval could be accomplished by relying on representations of the most recent list items in echoic memory, a short-lasting memory to which all spoken inputs have obligatory access (Crowder & Morton, 1969).

Hockey showed that when a passive strategy is adopted, increasing the rate of presentation improves recall accuracy, indicating reduced susceptibility to time-based decay known to characterise echoic memory (Hockey, 1973; see also, Bunting et al., 2006). In a computational model simulating such rapid presentation in running span, decay plays a crucial role in supporting successful recall (Weems, Winder, Bunting, & Reggia, 2009).

An active running span strategy involves attending to and processing incoming items while updating the target recall set so that only the relevant *n* items are maintained in WM. Pollack, Johnson and Knaff (1959) posited that participants continuously update after presentation of each item by dropping the old item from the target set and adding the new one. Postle and colleagues decomposed the potential stages involved in this updating process into five distinct but coordinated operations: encoding, discarding, repositioning, storing and rehearsing (Postle, 2003; Postle, Berger, Goldstein, Curtis, & D’Esposito, 2001).

A similar drop-and-capture process has been explored in tasks involving updating on the basis of semantic or spatial category (e.g. Kessler & Oberauer, 2014, 2015). Typically, in these tasks a single item (presented in a particular location or belonging to a specific category) is identified as irrelevant and has to be updated with a new item. Proposed mechanisms to support this updating process include item removal (Lewis-Peacock et al., 2018; Oberauer, 2018), inhibition of irrelevant items (Hasher & Zacks, 1988; Jonides, Smith, Marshuetz, Koeppe, & Reuter-Lorenz, 1998), and attentional shifts away from the no-longer relevant representations (Cowan, 2001; McElree, 2001). In each case, these accounts assume that updating only alters the to-be-updated item at each step, leaving the representations of the items and their contexts for the remaining items unchanged.

The distinctive feature of updating in running span is that the position of each item in the to-be-remembered portion of the sequence changes as a new item is added to the list. At each updating step, the first item must be discarded and the second item must become the first item, and so forth. Such repositioning is integral to running span updating (Postle et al., 2001; Postle, 2003). An extension of the item-removal model might enable such repositioning (Oberauer, Lewandowsky, Farrell, Jarrold, & Greaves, 2012). Serial order in this model is encoded as a context code associated with each item using Hebbian learning. It was proposed that item removal is supported by an opposite process of Hebbian anti-learning involving unbinding an item from its positional context (Lewis-Peacock et al., 2018; Oberauer, 2018). In running span, the no-longer-relevant item could be unbound to remove it from the target set, while the remaining items could be repositioned by successively unbinding them from their current positions and rebinding them to new positions in the list.

An alternative possibility is that updating in running span is supported by resetting the position at which recall commences rather than modifying all item-order associations with each successive update. For example, the primacy model of serial recall represents serial order as a noisy activation gradient that diminishes across successive list positions (Page & Norris, 1998). To shift the start position in updating epochs at positions *n*+1 and beyond, the first item in the list (with the highest activation) could be suppressed following its retrieval.

The activation gradient across the remaining items would preserve relative serial order, such that the second item in the encoded list would have the highest activation level and would therefore be retrieved first. This single, executively-mediated process of suppression would involve fewer cognitive operations than multiple unbindings and rebindings of item and order representations for the entire memory set of *n* items.

Active updating in running span is a time-consuming process. When participants were instructed to use an active strategy, their recall accuracy improved as the rate of presentation was slowed (Hockey, 1973). A benefit of slow presentation rate on recall emerged even in the absence of explicit strategy instructions (Bunting et al., 2006). This suggests that updating in running span is a slow process that cannot be efficiently applied when successive items are presented rapidly. Lengthening the inter-item interval appears to provide the required time for the process to occur and imparts a performance advantage (Bunting et al., 2006; Hockey, 1973).

Updating is also cognitively demanding. Self-report data suggest that participants find slow-paced running span associated with active updating to be more challenging than passive listening in the fast-paced task (Bunting et al., 2006). Morris and Jones (1990) found that performance accuracy was superior for short lists that required no updates of the target recall set compared with longer sequences that do require updates. Recall was independent of the number of updating steps. This indicates that the resources required to support updating are independently deployed and replenished within each inter-stimulus interval (1s duration). This suggested a resource recovery rate or refractory period associated with updating (Morris & Jones, 1990; see also, Postle et al., 2001).

This evidence indicates that serial updating in running span is a complex and cognitively demanding process that is sensitive to the temporal parameters of the task. With rapid presentation rates there is inadequate time to update and participants opt for passive listening. When updating does occur, it increases recall accuracy and consumes resources. With a sufficiently long interval between items, these depleted resources may be restored in time for the next updating epoch. In Experiment 1 the temporal dynamics of the updating process in a slow-paced running span are examined. Experiment 2 investigates the extent to which temporal signatures of updating identified in Experiment 1 are tied to the relatively slow presentation rates associated with active updating.

Both experiments assess the demand on general cognitive resources in running span using a divided attention methodology (Craik et al., 1996). The logic is simple: if two tasks are performed together and given equal priority they should compete for the same general cognitive resources. The resources available for each task should therefore diminish, generating a dual task cost (Craik, Govoni, Naveh-Benjamin, & Anderson,1996). A similar approach adopted by Thalmann et al. used a concurrent choice reaction time (CRT) task between stimulus presentation and recall to examine the cognitive demands of different maintenance strategies for verbal items (Thalmann, Souza, & Oberauer, 2019). In the present experiments, a continuous CRT task was applied concurrently with running span. Delays in responding in the CRT task yielded a reaction time function tracking the magnitude and time course of resource-demand characteristics specific to running span. Importantly, in the present set of experiments the concurrent task was applied during the presentation phase of the memory task, allowing us to use RT costs of time-locked events during stimulus presentation to identify the time course of the cognitive demands of updating.

## Experiment 1

It was anticipated that a slow rate of presentation would encourage the use of an active updating strategy in running span (Bunting et al., 2006; Hockey, 1973). The continuous demand metric from the CRT task allowed us to investigate whether this demand varied as a function of task events or remained constant over the course of the task. Cognitive resources were anticipated to only be required to support updating when the presented lists were longer than the target number of items (Morris & Jones, 1990; Postle et al., 2001). A heightened demand (detectable by increased RTs on the concurrent CRT task) was therefore expected to be associated with later items in the list (*n+1*th position onwards) compared with the preceding items. This recruitment of resources to support target updating was specifically predicted to follow encoding of each item in the sequence beyond the *n*th position.

The resource demands of running span were compared with those of two other serial recall tasks. The first was simple span, a standard serial recall task that does not require updating of the encoded memory items. If running span indeed requires an updating process, we predicted that it would impose a greater cognitive demand than simple span. An absence of any differential demand between running span and simple span would suggest the use of a passive strategy in the former. The second comparison task was a modified version of simple span. In this, the start position for serial recall within a lengthy sequence was repeatedly reset, but a continuous update of the memory set was not required. In this, lengthy memory sequences that contained intermittent signals were presented. Participants had to prepare to recall the fixed set of items following the most recent cue in the list. Thus, each cue flagged a new start position that enabled participants to begin encoding a fresh set of target items and disregard all items held in memory preceding the cue. This allowed us to distinguish the cost associated with the complete memory reset, as in modified span, with the updating process requiring maintenance of a partial target set, as in running span. Further, a comparison of simple and modified span allowed us to assess if there is any discernible cognitive cost to a simple reset.

## Method

### Participants

Ninety-two native English speakers were included in the study. Complete data were recorded for 90 participants (68 female, mean age= 24.38 years, SD= 4.04 years). Participants were recruited using printed and electronic advertisements within and beyond the MRC Cognition and Brain Science’s research participation system. Informed consent was obtained in accordance with ethical approval from Cambridge Psychology Research Ethics Committee (PRE 2016.066), and participants were compensated for their time and travel costs.

### Design

The study used a 3×2 mixed factorial design. Task was a between-group factor; participants were randomly assigned to one of three groups completing the different memory tasks (simple span, modified span, and running span). Attentional load was a within-subject factor, with each participant completing two load conditions (single and dual task). In addition, all participants also completed a digit span task for an assessment of verbal WM capacity.

### WM tasks

Three WM tasks (Figure 1A) were completed by separate groups of participants. The task instructions, length of lists, and number of target items varied across tasks as detailed later. The size of the target recall set within each task was determined on the basis of pilot data; the WM load associated with an accuracy between 75-85% in each memory task respectively was selected. Recall attempts were digitally recorded using a microphone on a headset and subsequently transcribed. Recall accuracy was measured in terms of the proportion of items recalled in the correct serial position.

**Figure 1.**
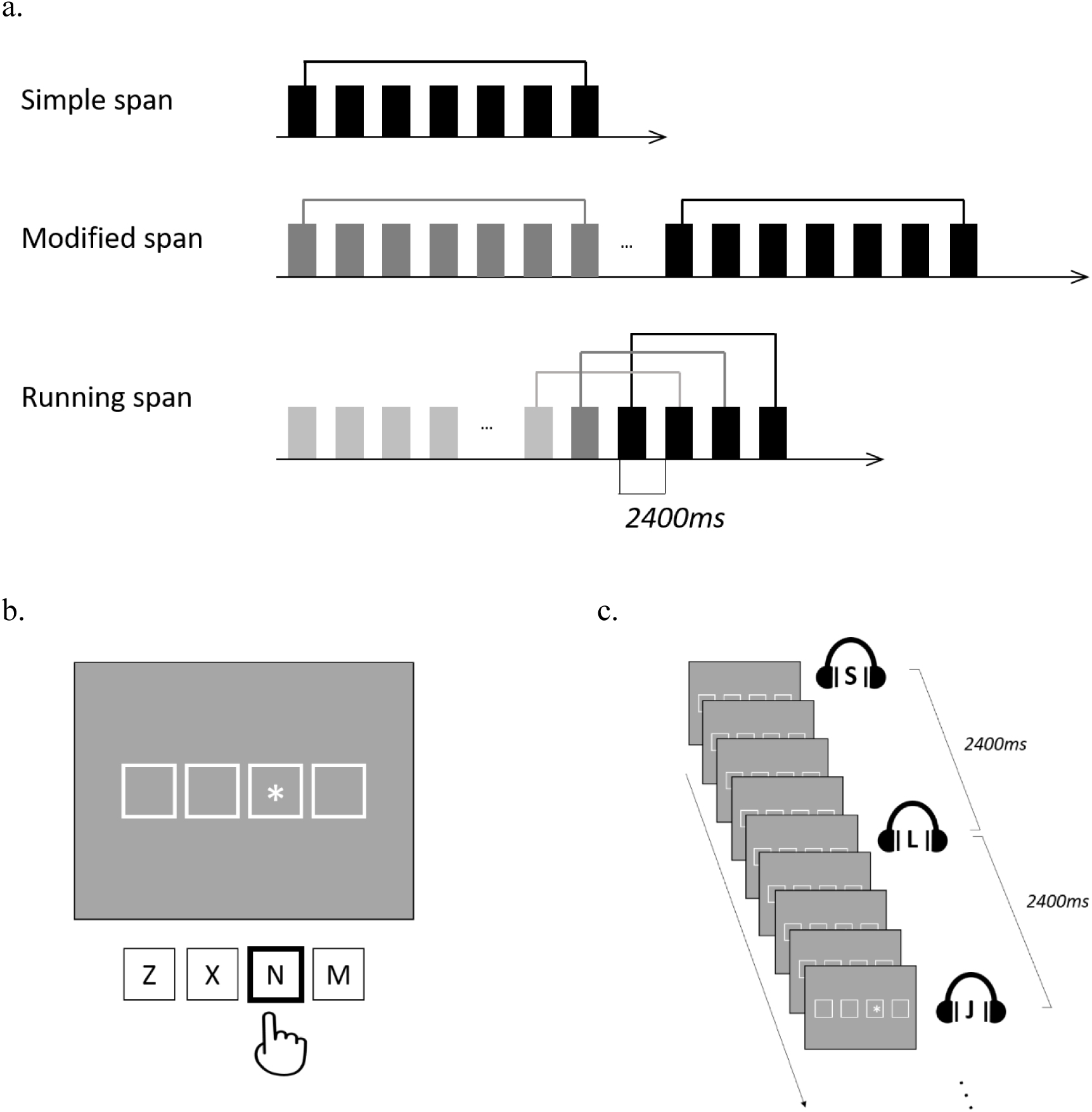
Task design. a. Schematic of a memory list for each working memory task. Memory items (marked in rectangular boxes) were presented sequentially at a rate of 2400ms per item (800ms for item, followed by 1600ms of silent interval) using spoken presentation. List length varied in modified span and running span. The later items in the list were relevant for recall (marked in black, varied as per task), while earlier items were not (grey). The brackets above items denote the items in the same target recall set at a given timepoint. b. The continuous reaction time (CRT) task. Four square frames were presented on screen corresponding to four response keys. Participants pressed the key corresponding to the frame containing the star. A CRT stimulus was presented immediately following the response to the previous stimulus. c. Dual task structure with a simultaneous application of (auditory) working memory + (visual) CRT task. Memory items presented every 2400ms; CRT task was participant paced, so the number of CRT stimuli presented between each memory item varied contingent on participant reaction times.

### Running span

Participants attempted to recall the last four items of the presented list in serial order. Lists started with a 1s tone and contained 4 to 12 items. The length of the list was unknown to the participants. Each block contained 10 trials, including 1 presentation of each list length, except for lists containing 8 items that were presented twice in a pseudo-random order. Participants thus completed fifty trials over five blocks.

### Modified span

Participants heard sequences of letters periodically interspaced with cues that indicated the start of an ordered list of items they were to remember. Lists contained 7, 14, 21, or 28 items, and the length of each upcoming list was unknown to participants. The cue was a 1s tone, presented at the start of each list, and after every 7^th^ item in the case of longer lists. Participants were told to recall the items in serial order following the latest tone. Irrespective of list length, the number of items to be recalled was always fixed at seven, as in simple span. Each list length was presented twice in a pseudo-random order in a block. There were forty trials presented over five blocks.

### Simple span

Each list started with a 1s tone and contained seven items. Participants attempted to recall all list items in the original serial order. Fifty lists were presented over five blocks.

Identical protocols were employed to present stimuli and generate sequences in all three memory tasks. Lists were generated randomly subject to the following constraints: (a) 20 consonants were used as stimuli (‘w’ was excluded), (b) a letter could only be repeated after every 7 items, (c) two phonologically similar letters could not be presented consecutively, (d) three or more letters could not be presented in alphabetic order. The letters were spoken by a male speaker of British English and were recorded at a sample rate of 44.1 kHz. Sound files containing each letter were edited to be 800ms long and the location of the letter was adjusted so that letters sounded evenly spaced. (Morton, Marcus, & Frankish, 1976). The tasks used sequential auditory presentation over headphones (Sennheiser HD 280 PRO II) at a rate of 2400ms per item (i.e., 800ms for item presentation followed by a silent interval of 1600ms). Each list was preceded by a 1s tone to alert participants to the start of the trial and a fixation cross was displayed on the screen to maintain fixation while attending to the auditory stimuli. The duration of the presentation phase varied with list length and was always immediately followed by spoken serial recall for a maximum of 20s.

### Continuous reaction time (CRT) task

In this task (Figure 1B), an asterisk was presented in one of four possible square frames on the screen, and participants were required to press the key corresponding to the frame containing the asterisk as quickly and accurately as possible. Following a key-press the asterisk immediately shifted to one of the other three frames chosen at random. Responses in this task involved key-presses, using the first two fingers of the right and left hands. The task was self-paced, such that the onset of the next stimulus immediately followed each response. Reaction time and accuracy were both recorded in this task.

### Dual task condition

In the dual task conditions (Figure 1C), participants simultaneously performed the WM and CRT tasks and were instructed treat each task with equal priority. The presentation protocol and list and stimulus generation in the dual task conditions were identical to their corresponding single task conditions. The visual presentation of the first stimulus in the CRT task was synchronized with the auditory onset of the memory sequence in the respective WM task. Thereafter, the CRT stimuli were presented successively during the presentation of the memory list, and ceased when the retrieval phase of the WM task started. The aim in the dual task was to respond in the CRT task as quickly and accurately as possible using keypresses, while also attending to the memory list for subsequent recall in the respective WM task.

In both single and dual CRT task, only responses occurring at least 200ms after CRT stimulus onset were recorded to capture psychological processes rather than anticipatory motor responding (Van Zandt, & Townsend, 2014). CRT responses due to accidental holding down of a response key from the previous trial were excluded from analysis. Also, programming constraints caused the CRT task to abruptly stop upon reaching the end of the memory list in the dual task condition, which truncated the recording of any CRT responses that may have followed. To remove the possibility of bias, CRT responses associated with the final item in each memory list across running, modified and simple span were excluded from analyses.

### Digit span task

In addition to the tasks above, a digit span task was administered to measure verbal WM capacity. For this, the digits 0 through to 9 were presented sequentially in the centre of the screen for 1000ms followed by a blank inter-stimulus interval (ISI) for 1000ms. Stimuli were pseudo-randomly selected such that stimulus repeats were only allowed after every nine items. At the end of the list, a 3×3 grid was presented on screen and participants recalled the presented sequence in serial order by indicating responses with mouse clicks. The task commenced with lists containing 4 items. At each span level, 4 trials were presented and participants advanced to the next level (i.e. the list lengthened by 1 item) if they responded with 75% accuracy at any given level. The task ended once performance failed to meet this condition. Span was recorded as the level at which participants correctly recalled 3 or more trials.

### Procedure

For each participant, the digit span task was administered first, followed by three experimental tasks: the continuous reaction time (CRT) task, the respective working memory (WM) task, and the dual task condition in which both tasks were performed concurrently. These were implemented using a blocked design with a fixed task order (CRT, WM, and dual task) repeated over 1 practice block and 5 experimental blocks. All tasks were completed within one session, typically lasting between 1.5 to 2 hours. All tasks were designed and presented on a PC using MATLAB 2014a (The Mathworks, Inc.) and Psychtoolbox-3 (Brainard, 1997; Pelli, 1997; Kleiner et al, 2007).

## Results

There were no group differences in age, *F*(2,87) = 2.14, *p* > .05, gender, *χ^2^*(2, N = 90) = 0.12, *p* > .05, or verbal WM capacity measured using the digit span task, *F*(2,87) = 0.21, *p >* .05. Mean recall accuracy and RTs from the WM and CRT tasks respectively are presented in Table 1 for both load conditions across the three WM tasks.

**Table 1.** Mean ± SDs for performance in concurrent choice reaction time (RT) task and primary memory task, for each load and task condition

|  |  | Running Span | Modified Span | Simple Span |
| --- | --- | --- | --- | --- |
| RT (ms) | Single | 443 ± 66 | 436 ± 56 | 437 ± 46 |
| | Dual | 540 ± 143 | 499 ± 77 | 474 ± 110 |
| Recall accuracy<sup>1</sup> | Single | .83 ± .10 | .70 ± .21 | .77 ± .15 |
| | Dual | .76 ± .13 | .60 ± .24 | .67 ± .16 |

<sup>1</sup> Recall scored as proportion of items recalled in correct serial position

### Reaction time data

The delay in the remaining CRTs was analysed at three levels: as a function of overall WM task, events within trials (serial position and item types), and latency from item onset.

#### Task level

RTs in the single-and dual-CRT task were compared across the three groups completing different concurrent memory tasks using a 2×3 mixed measures ANOVA (Figure 2). Responses in the single CRT task were faster than in the dual task condition, *F*(1,87) = 51.57, *p* < .05, 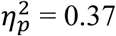. A significant interaction between load and memory task was observed, *F*(2,87) = 3.60, *p* < .05, 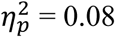. To understand this interaction, a simple one-way ANOVA was conducted comparing the dual task RT cost (computed by subtracting single CRTs from dual CRTs) across the three memory tasks. Results revealed that the dual task cost was greater during concurrent execution of running span than that during simple span, mean RT difference = 60ms, *p* < .05. Dual task cost in RTs during modified span did not differ from that during running or simple span, *p* > .05 in both cases.

**Figure 2.**
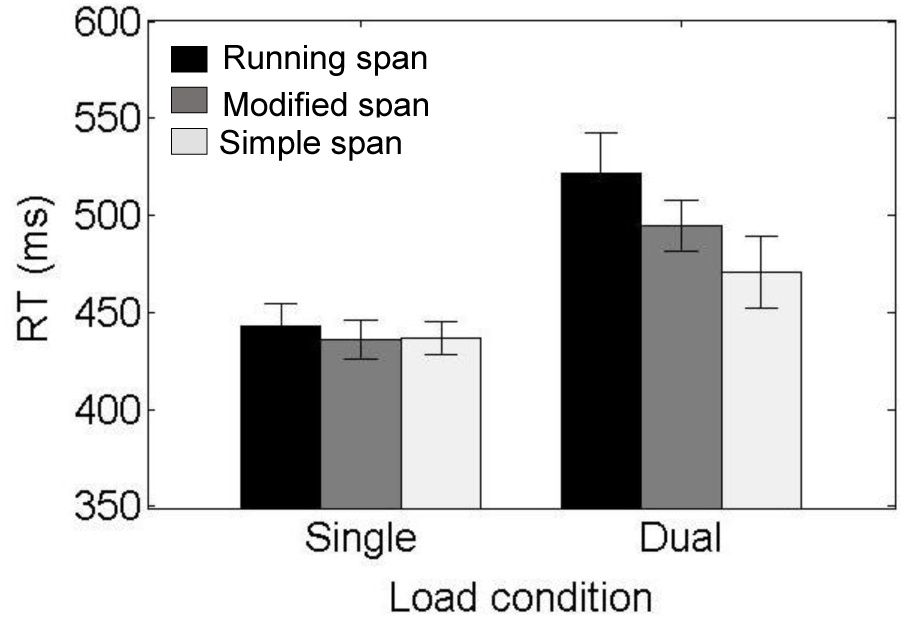
Mean single and dual RTs from the concurrent CRT task across the groups undertaking different memory tasks. Error bars represent standard error of the mean.

#### Trial level

To trace the temporal source of this differential demand (Figure 3a), trials were divided into early (positions 1-4) and late (positions 5-7) serial positions. For comparability across the three tasks, only the first 7 serial positions from each memory sequence were included in this analysis. RTs were segmented into their respective serial positions and compared across tasks using one-way ANOVAs. For the early positions in the trial, no difference in RTs was found across tasks, *F*(2,87) = .72, *p* > .05. For later positions, a significant task effect was observed, *F*(2,87) = 3.64, *p* < .05, such that RTs associated with these later positions were longer in running span relative to simple span (mean difference = 118ms, *p* = .05). The same positions in modified span did not differ significantly from either simple or running span, both *p*s > .05.

**Figure 3.**
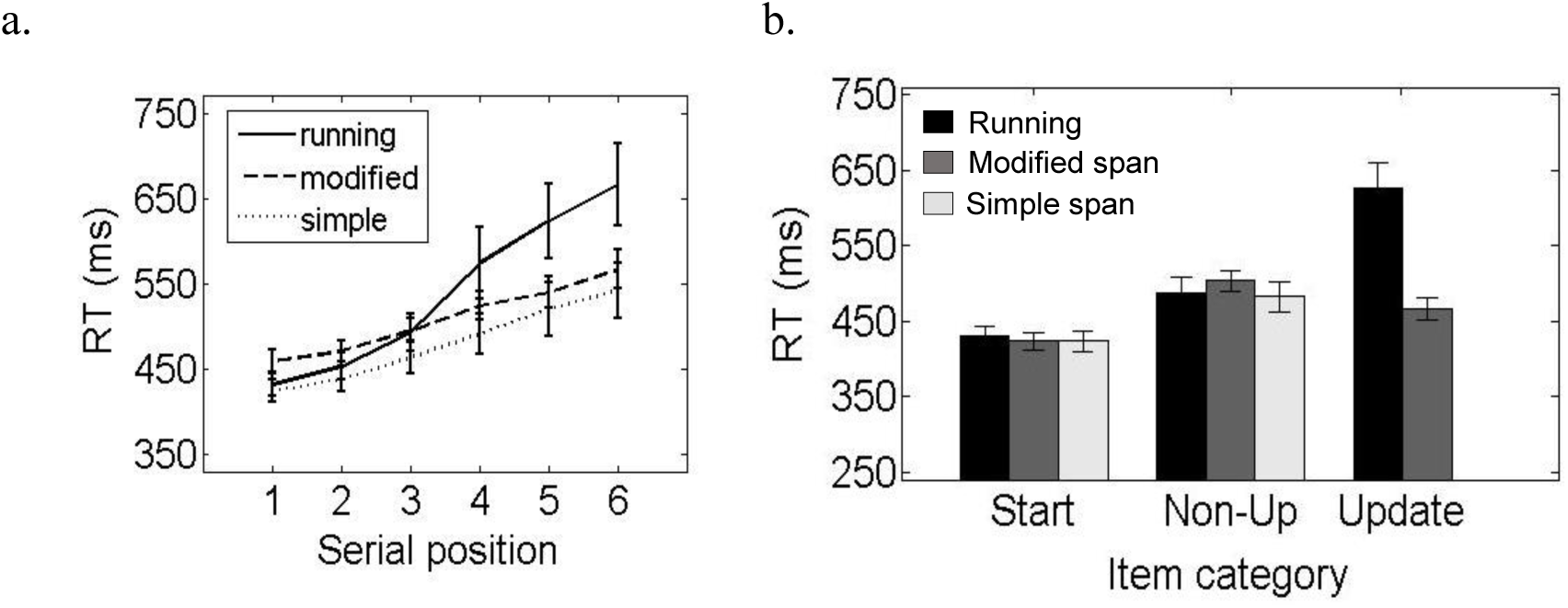
a. Concurrent RTs across six serial positions in the list for each memory task. RTs associated with the final position in the list (=7 for simple span) were excluded from analysis and thus not displayed here, see text for data exclusion. b. Concurrent RTs as a function of category of item category for all three memory tasks. See Table 2 for item classification details; simple span did not have any update items. Error bars represent standard error of the mean.

While the above analysis allowed comparability across tasks, it excluded data from later positions in running span lists. To include all positions, items were classified into three categories – start, non-update, and update items (detailed classification provided in Table 2). Concurrent RTs were tagged as one of the three item-types and compared using one-way ANOVAs across the three WM tasks (Figure 3b). No difference was found in RTs across the three tasks during both start items, *F*(2,87) = 0.06, *p* > .05, and non-update items, *F*(2,87) = 0.30, *p* > .05. Further, RTs associated with update items were significantly slower in running span than in modified span, *t*(58) = 4.07, *p* < .05, mean RT difference = 203ms, Cohen’s *d* = 1.05.

**Table 2.**
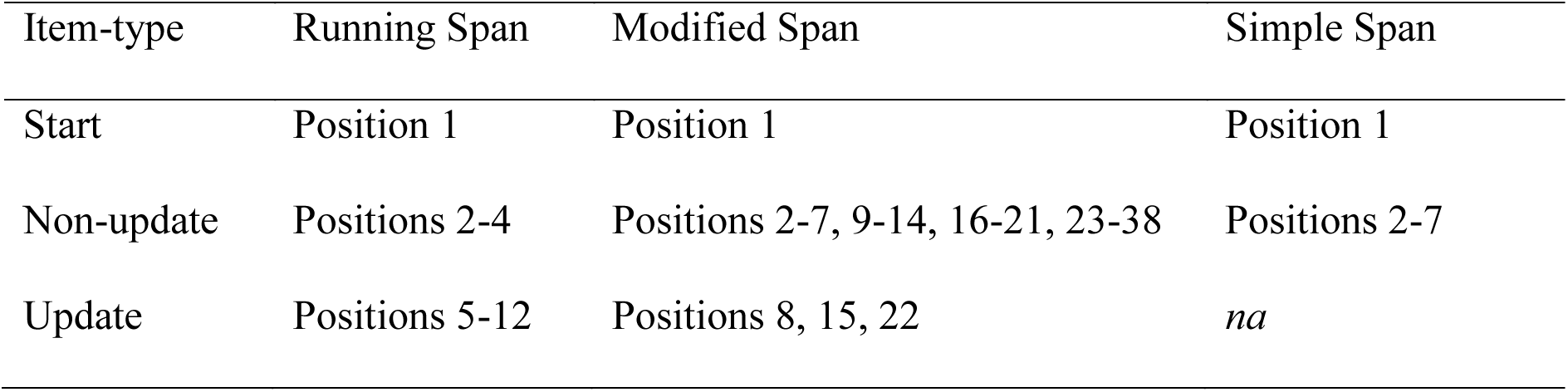
Detailed classification of serial position into item categories for each memory task

#### Item level

RTs were also analysed as a function of latency from the onset of memory item. The inter-item duration of 2400ms (including 800ms of stimulus presentation and 1600ms of silent ISI) was divided into six micro intervals of 400ms each. Stimulus-locked RTs from the concurrent task were segmented into these micro intervals and compared using one-way ANOVAs (a) across items within running span, and (b) between running and modified span (Figure 4).

**Figure 4.**
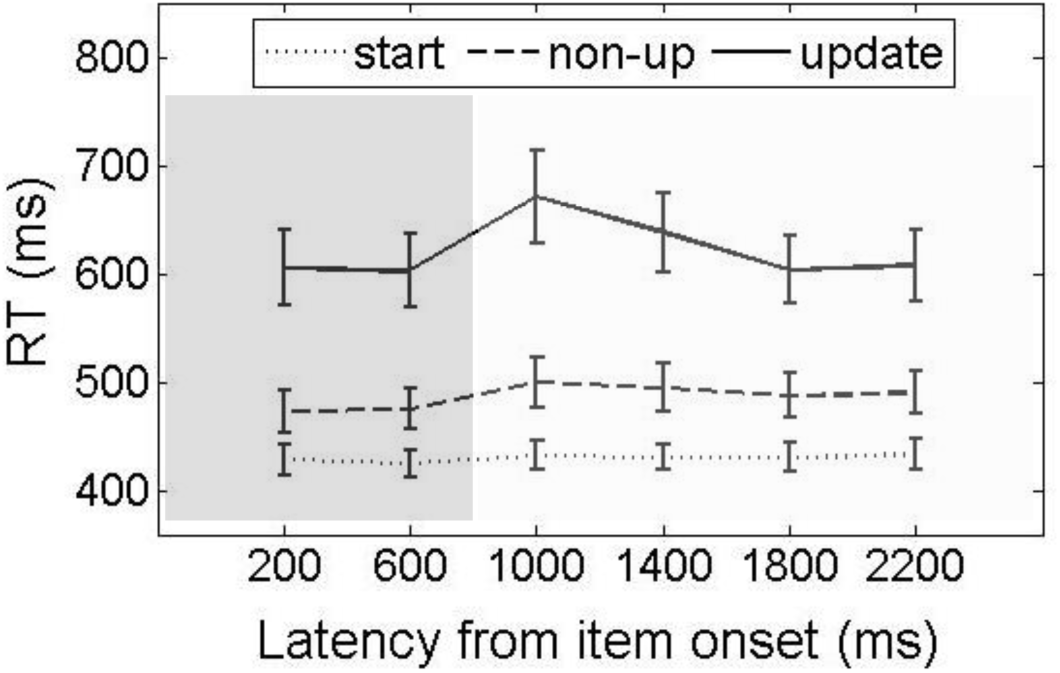
Concurrent RTs as a function of latency from item onset of memory item in running span, separated by item-type. The first 800ms represent the duration of the item presentation (shaded in dark grey), followed by a 1600ms silent inter-item interval (shaded in light grey). Error bars represent standard error of the mean.

Update items in running span showed a significant effect of interval, *F*(5,145) = 6.84, *p* < .05, 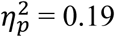. Pairwise Bonferroni comparisons revealed that for updating items, responses were at a stable level from the first to the second interval, and then showed a sudden slowing in RT from the second to the third interval, mean RT difference = 92ms, *p* < .05. From the third interval onwards responses gradually returned to the baseline level with non-significant increments in RTs, all *p*s > .05. Non-update items in running span also showed an interval effect, *F*(5,145) = 3.42, *p* < .05, 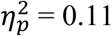. This was driven by a small difference between the first and fourth interval, mean RT difference = 22ms. All other pairwise comparisons of the micro-intervals within non-update items in running span were non-significant. The start items in running span showed no interval effect on RTs, *F*(5,145) = .74, *p* > .05. Update items in running span were distinguished by a peak delay in the third interval (around 1000ms), while those in modified span showed no effect of interval, *F*(5,145) = 1.36, *p* > .05.

To ensure that the results reported here were not due to any data outliers, the analyses were repeated with the following trimming and outlier correction procedures. First, RTs associated with inaccurate CRT responses were excluded. Second, RTs outside of individual mean ± 2.5 standard deviations were removed. Finally, participants whose mean RTs exceeded the group mean RT by ± 3 standard deviations were removed (one participant removed from the running span group, and one from the modified span group). These exclusions did not lead to any change in the pattern of results.

## Recall data

Recall performance across serial positions was analysed separately for each task using repeated measures ANOVAs. In running span, recall accuracy was higher in single-than in dual-task load, *F*(1,29) = 22.19, *p* < .05, 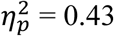. Across both load conditions, recall improved with each consecutive serial position, *F*(3,87) = 121.23, *p* < .05, 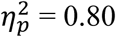. The impact of load interacted with serial position, *F*(3,87) = 19.68, *p* < .05, 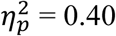, with a decreasing effect of load on recall accuracy for each consecutive serial position, from *d* = 1.16 (position 1), = .76 (position 2), = .52 (position 3), and = .19 (position 4). Pairwise Bonferroni comparisons indicated that the difference between single and dual task load was significant for serial positions 1-3, *p*s < .05, but not for the final position, *p* = .33.

Recall in modified span exhibited a significant effect of both load, *F*(1,29) = 47.59, *p* < .05, 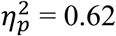, and serial position, *F*(6,174) = 53.82, *p* < .05, 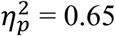. Load and serial position showed a significant interaction, *F*(6,174) = 3.55, *p* < .05, 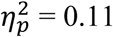, such that the effect of load on recall increased from serial position 1, *d* = .89, to position 4, *d* = 1.41, and then subsequently decreased from position 5, *d* = .98, to the final position, *d* = .44., all *p*s < .05.

Recall in simple span showed a significant effect of load, *F*(1,29) = 53.29, *p* < .05, 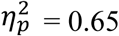, and serial position, *F*(6,174) = 57.03, *p* < .05, 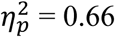. Load and serial position showed a significant interaction, *F*(6,174) = 8.61, *p* < .05, 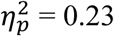, such that the effect of load on recall increased from serial position 1, *d* = .76, to position 5, *d* = 1.58, and then subsequently decreased from position 6, *d* = 1.32, to the final position, *d* = .45, all *p*s < .05.

In summary, a strong dual task cost in recall was observed in all three tasks. However, serial position functions differed (Figure 5): recall accuracy in running span improved with each consecutive serial position, whereas performance in both modified span and simple span decreased consecutively until position 4-5 and then subsequently improved for the remaining positions.

**Figure 5.**
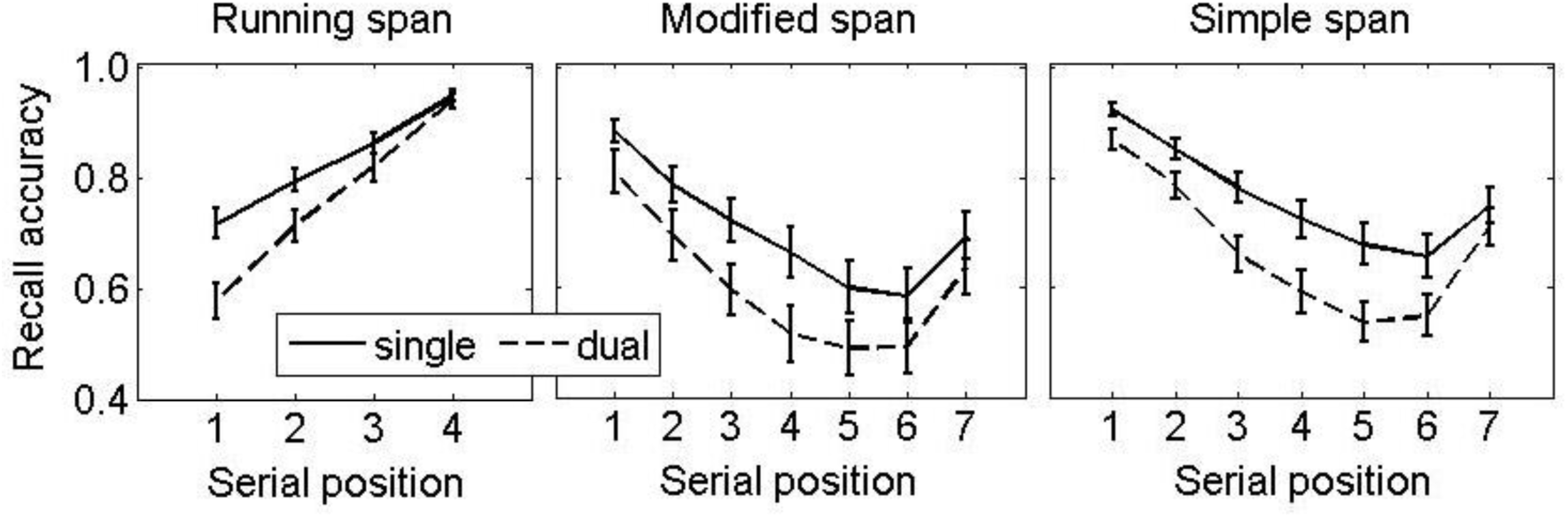
Recall accuracy (proportion of items recalled in correct serial position) across each serial position for the three memory tasks across single (solid line) and dual (dashed line) load conditions. Error bars represent standard error.

## Discussion

Concurrent RTs were slower when participants simultaneously performed running span relative to both simple and modified span conditions. In running span positions following the *n*th position (item 4) generated a substantial RT cost with an additional peak approximately 1000ms following the onset of these later items. This diminished to baseline level over the course of the item interval, prior to the next item presentation.

These data reinforce previous claims that the cognitive demands associated with running span reflect the use of an active strategy (e.g., Bunting et al., 2006; Hockey, 1973; Morris & Jones, 1990, Postle, 2003). The data suggest the initiation of an updating mode that imposes a considerable demand on resources during the list positions beyond *n*. The specificity of the updating-related resource demand to the positions beyond *n* was consistent with the hypothesis advanced by Morris and Jones (1990; see also Postle et al., 2001) that an updating operation is only set into action when a memory sequence contains *n* items.

In the interval between two updating items in running span, a sharp demand was observed at 1000ms from item onset which returned to baseline level prior to presentation of the next memory item. This suggests a process that updates the target recall set when prompted by a new item, rather than a generalized state enabling constant intervention. It is consistent with a previous suggestion that individual episodes of updating dos not impose cumulative demands on cognitive resources (Morris & Jones, 1990; Postle et al., 2001). The time course of initiating and completing the complex updating process underlying this peak could be fixed or could be deployed flexibly according to task conditions, such as the rate of presentation employed. The data from this study do not allow us to differentiate between these alternatives.

The externally-signalled memory reset event in modified span placed little demand on cognitive resources compared with the substantial cost of an internally-driven, serial update of already encoded items in running span. Both the concurrent RT and recall data from modified span corresponded closely to simple span, a standard serial recall task with no embedded target updates. The process of starting encoding afresh mid-sequence therefore appeared to be equivalent to starting at the beginning of a list.

If the 1000ms peak demand reflects updating, it should diminish or even disappear in conditions that do not favor an active strategy. We tested this prediction next by manipulating the opportunity for participants to use an active strategy. Previous investigations showed that when faced with a faster rate of presentation, participants shift to passive listening, a strategy that demands few resources during item presentation (Bunting et al., 2006; Cowan et al., 2005; Weems et al., 2009). It was therefore predicted that fast presentation rates would reduce active updating and decrease the demand associated with the running span task.

## Experiment 2

Previous studies suggest that a passive strategy in running span is favored when the rate of presentation is faster two or more items per second while active updating is employed with rates slower than one item per second (Bunting et al., 2006; Cowan et al., 2005; Hockey, 1973; Broadway & Engle, 2010; Botto, Basso, Ferrari, & Palladino, 2014; Bunting et al., 2006; Collette et al., 2007; Elosua & Ruiz, 2008; Kiss et al., 1998; Morris and Jones, 1990; Postle, 2003; Postle et al., 2001). Results from Hockey (1973, see also Hamilton & Hockey, 1974) show that in the intermediate interval the performance benefits associated with active updating progressively increase at the rate slows, indicating that during this period participants may have a choice between the two strategies or may even mix them across trials.

In Experiment 2 three presentation rates were used – fast (400ms/item), medium (800ms/item) and slow (1600ms/item). Each running span condition was combined with the concurrent CRT task, as in Experiment 1. It was anticipated that the resource demand indexed by the CRT cost would increase as the presentation rate slowed as a consequence of a shift from a passive listening to active updating. The intermediate presentation provided the opportunity to examine the boundary conditions for deployment of an active processing strategy. It also allowed us to determine whether the demand on resources associated with has a fixed time course or shifts according to presentation rate. If it is flexibly applied, the resource burst detected at 1000ms after item presentation with the slow presentation rate (2400ms) in Experiment 1 would be expected to move forward in time in even the slowest rate of 1600ms in Experiment 2.

## Method

### Participants

Thirty participants (18 female, mean age = 24.3 years, SD = 3.9 years) who spoke English as a native language were recruited for this experiment. The total number of participants tested in this experiment was less than in the first experiment, but per condition both experiments tested the same number. In the first experiment, 30 participants completed one of three conditions in a between-group design. In this experiment, the same 30 participants completed each of the three conditions in a within-subject design. The experiment was granted ethical approval by the Cambridge Psychology Research Ethics Committee (PRE 2016.066), and participants were compensated for their time and travel costs.

### Design

This experiment used a 3×2 within-subject design to investigate two factors: presentation rate and attentional load. Participants completed running span with different rates of item presentation (fast, intermediate, and slow), counterbalanced across sessions. Attentional load was also manipulated, as performance was measured in both single and dual task conditions in each session.

### Task

The structure of running span task employed in this experiment differed from that used in Experiment 1 in two aspects.

First, a new set of stimuli were recorded so that the presentation of the memory items (i.e. letters) lasted for 400ms. For this, letters spoken in a single stream at a rate of 2 letters per second by a native British English female speaker were recorded. Using the P-centre approach (Morton et al., 1976), the audio was segmented into constituent letters and then compressed into 400ms files using Adobe Audition 3.0.

Second, the inter-item interval in the task varied across the rate conditions. The fast paced task involved successive presentations of items for 400ms with no intervals between them. In the mid-rate condition, a silent 400ms interval was interleaved between successive items, such that items were presented at a rate of 800ms/item. The silent interval was increased to 1200ms in the slow-rate condition presenting items at 1600ms/item. All other features of the task including modality of presentation and recall, list generation protocol, and number of trials and blocks in each session were the same as in Experiment 1.

### Strategy Data

At the end of each session, participants were provided with a list of six possible strategies and asked to rate the frequency with which they used each one on a 4 point scale ranging from almost never to almost always (Table 3). Further, at the end of the final session, participants were asked to report the rate condition they experienced as the least challenging.

**Table 3.**
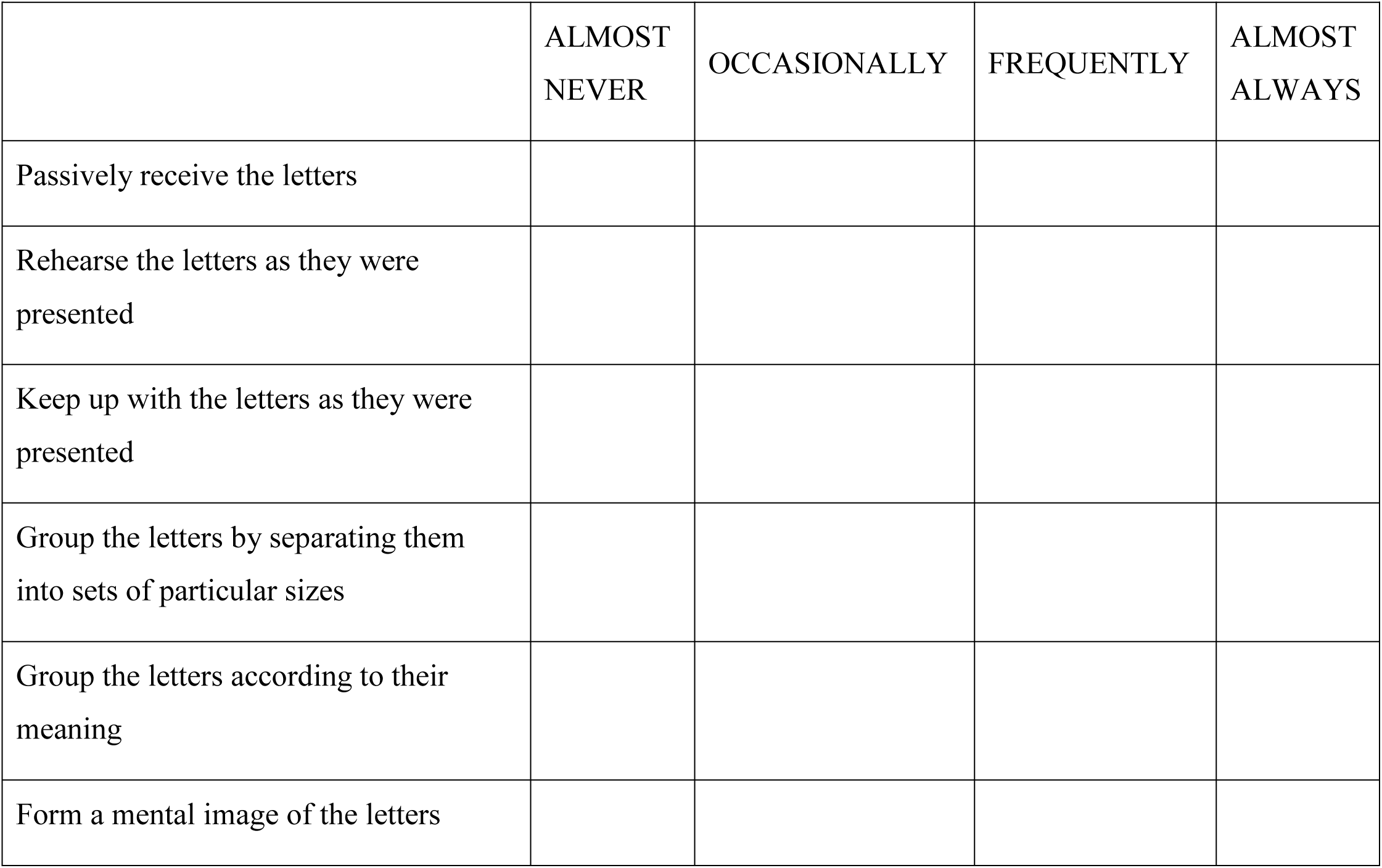
Strategy questionnaire completed after each rate condition In this session, while the list was being presented, to what extent did you...

### Procedure

Participants attended three sessions in this experiment. In each session, they completed three tasks: the continuous reaction time (CRT) task, the running span task with the respective rate-condition, and the dual task condition in which both tasks were performed concurrently. Task order was fixed (CRT, running span, and dual task) in each block and an initial practice block was followed by five experimental blocks. Each session concluded with the administration of the strategy questionnaire and lasted approximately an hour.

## Results

Mean recall accuracy and RTs from the running span and CRT tasks are presented in Table 3 for both load conditions across the three presentation rates.

### Reaction time data

RTs from the concurrent CRT task were analysed at three temporal levels, mirroring the investigation conducted in Experiment 1.

#### Task level

RTs in the single and dual CRT task were compared across the three rate conditions in a 2×3 repeated measures ANOVA (Figure 6). There was no effect of load, *F*(1,29) = .008, *p* > .05, a significant effect of rate, *F*(2,58) = 3.27, *p* = . 05, 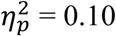 and a strong interaction between load and rate, *F*(2,58) = 40.79, *p* < .05, 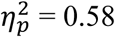. Pairwise Bonferroni comparisons examining this interaction revealed a significant dual task cost in the fast rate condition, *t*(29) = 4.74, *p* < .05, *d* = 1.05 and the slow rate condition, *t*(29) = 3.30, *p* < .05, *d* = .59. These costs were found in opposite directions: compared with the respective single RTs, dual RTs were faster in the fast paced condition, mean RT difference = 16ms, and slower in the slow-rate condition, mean RT difference = 19ms. RTs in the mid-rate did not differ significantly between the two load conditions, *t*(29) = .77, *p* > .05. Therefore, the demand placed on cognitive resources (indexed by the concurrent RTs) was greatest for the slow-rate condition and decreased with an increase in presentation rate.

**Figure 6.**
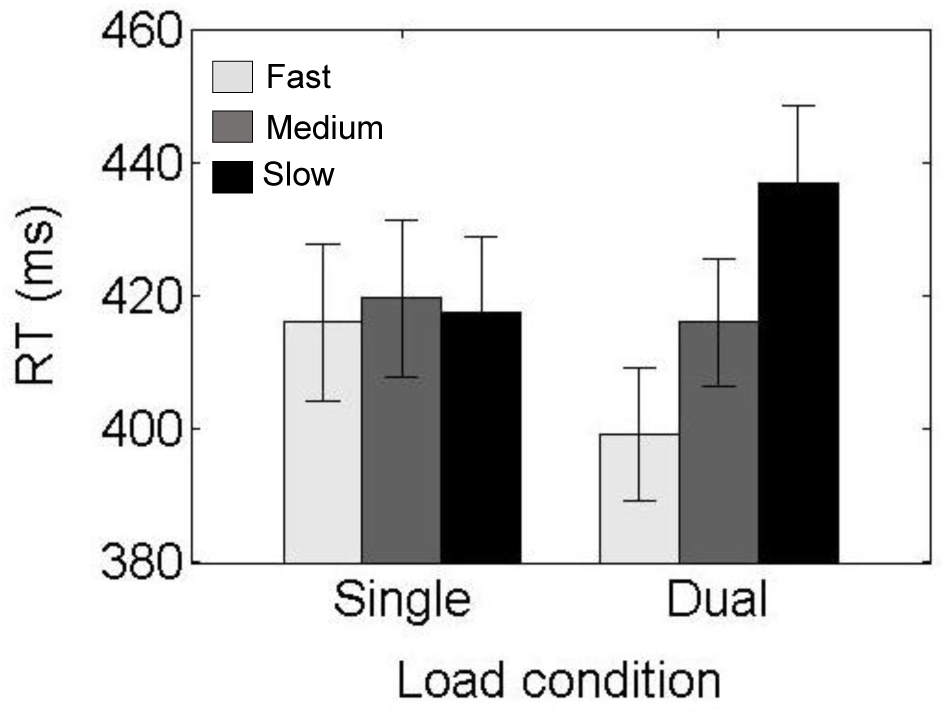
Mean single and dual RTs from the concurrent CRT task across three rate conditions. Error bars represent standard error of the mean.

#### Trial level

To examine whether this differential demand on resources found at the task level varied within each trial, we compared RTs across 11 serial positions for each of the 3 rate conditions (Figure 7). The 11×3 repeated measures ANOVA revealed a significant interaction between position and rate, *F*(20,560) = 4.89, *p* < .05, 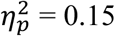, which was explored using separate analyses for each rate condition. For the fast paced task, pairwise comparisons of concurrent RTs between consecutive serial positions were not significant, all *p*s > .05. In the medium condition, RTs showed a gradual but non-significant increase at each consecutive position from 1 to 4, *p*s > .05, which was followed by a sharp rise in RTs from position 4 to 5, mean difference = 22ms, *p* < .05. This increased further from position 5 to 6, mean difference = 15ms, *p* = .05, and then stabilised from position 6 onwards, all *p*s > .05. In the slow paced task, RTs increased from position 1 to 5, mean difference between consecutive positions = [11ms, 14ms, 28ms, 19ms], all *p*s < .05. From position 5 onwards in the slow lists RTs were maintained at a steady level with no significant difference between consecutive positions, all *p*s > .05.

**Figure 7.**
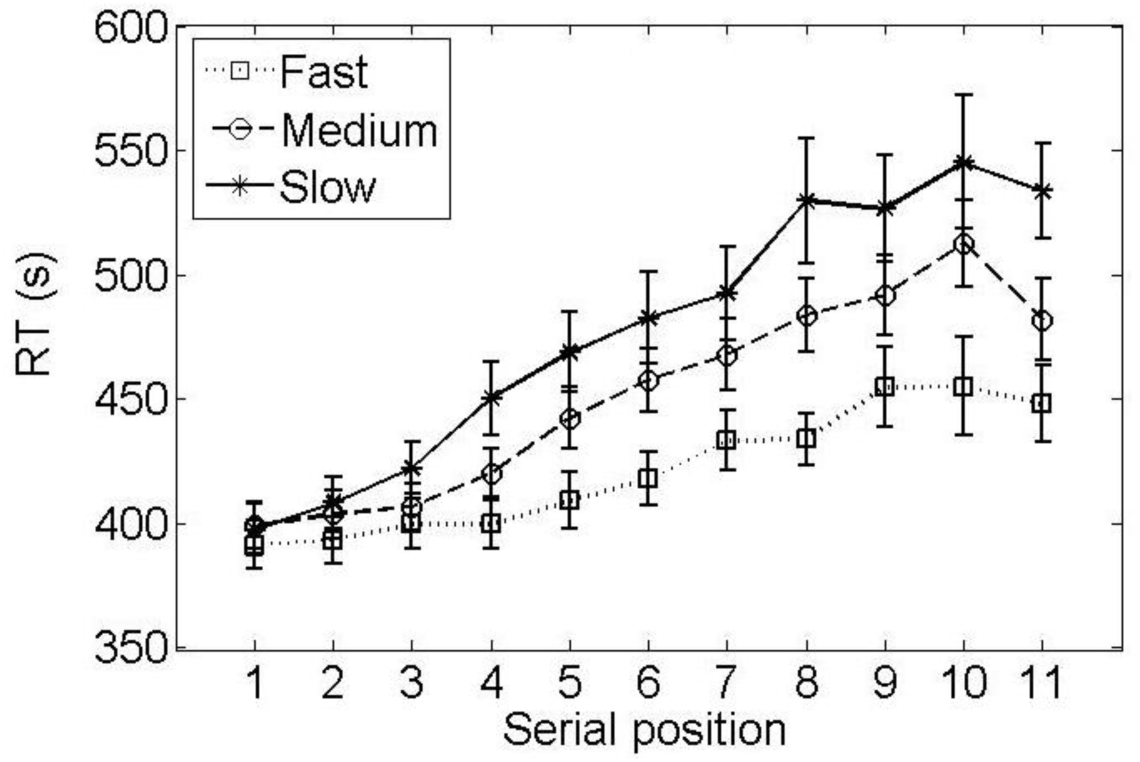
Concurrent RTs across serial positions for each running span condition. RTs associated with the final position in the list (e.g. position 12 for the longest list) were excluded from analysis and thus not displayed here, see text for data exclusion. Error bars represent standard error of the mean.

To trace the serial position at which these demand profiles diverged, the RTs associated with each position were compared across the rate conditions. The three rates did not differ significantly across the first two positions, *p*s > .05. At the third position in the list, RTs in the slow task were more delayed than the fast task, mean difference = 22ms, *p* < .05, a difference that increased over subsequent positions, mean difference > 51ms, *p*s < .05. At the fourth position, RTs in the medium rate condition also diverged from those in the fast paced task, mean difference = 21ms, *p* < .05. This significant difference was maintained over most of the subsequent positions, mean difference > 33ms, *p*s < .05, except at position 9 and 11, *p* > .05. The RTs in the mid and slow rate conditions were not different from each other across most positions, *p*s > .05, except position 4 and 11, mean difference = 30ms, = 53ms, respectively, *p*s < .05.

#### Item level

RTs were investigated as a function of latency from item onset from positions 5 onwards in the list (Figure 8). As in Experiment 1, the inter-item interval duration was divided into micro-intervals of 400ms each. In the medium rate condition, the total duration of 800ms between two consecutive item presentations (including 400ms of stimulus presentation and 400ms of silent ISI) was divided into two micro intervals. For the slow rate condition, a similar segmentation of the 1600ms inter-item duration resulted in four micro intervals, including one presentation and three post-presentation silent intervals. RTs were then classified into respective micro intervals for the slow and medium tasks. The fast condition was not included as the inter-item interval of 400ms could not be divided into further micro intervals. For the mid-rate condition, there was no difference in RTs between the two micro intervals for update items, *t*(29) = .89, *p* > .05. In the slow task, a significant difference between the micro intervals was observed for update items, *F*(3,87) = 13.19, *p* < . 05, 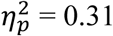. Pairwise Bonferroni comparisons revealed a delay in RTs in the third micro interval (around 1000ms post onset) compared with the other intervals, mean difference > 24ms, *p*s < .05.

**Figure 8.**
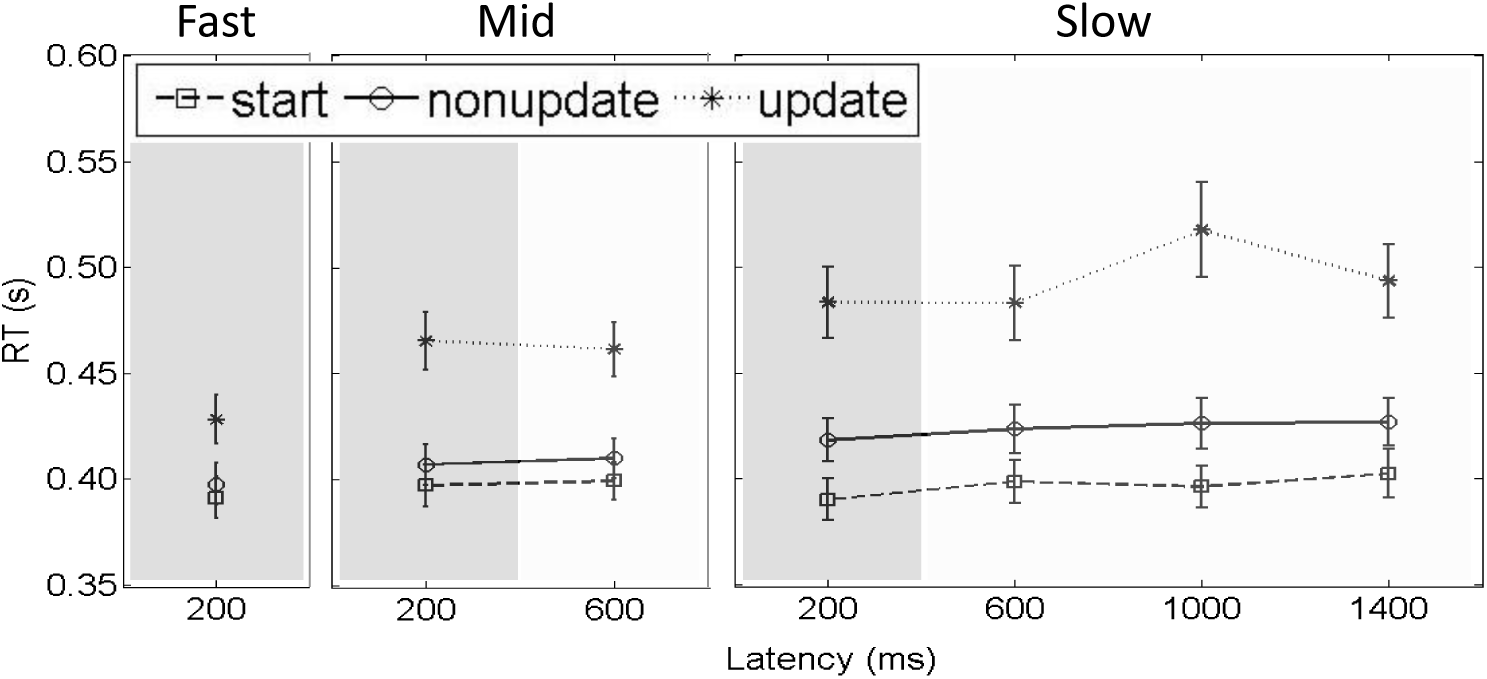
Concurrent RTs as a function of latency from item onset of memory item for all three rate conditions, separated by item-type. The first 400ms represent the duration of the item presentation (darker shading), followed by variable duration of silent inter-item interval (lighter shading). Error bars represent standard error of the mean.

As in Experiment 1, analyses were repeated to ensure that the results were not due to any outliers in the data. Removal of RTs associated with inaccurate CRT responses, RTs outside of individual mean ± 2.5 standard deviations, and RTs exceeding the group mean by ± 3 standard deviations in any rate condition were removed (one participant removed, N=29 analysed) did not lead to any change in the pattern of results.

**Table 4.**
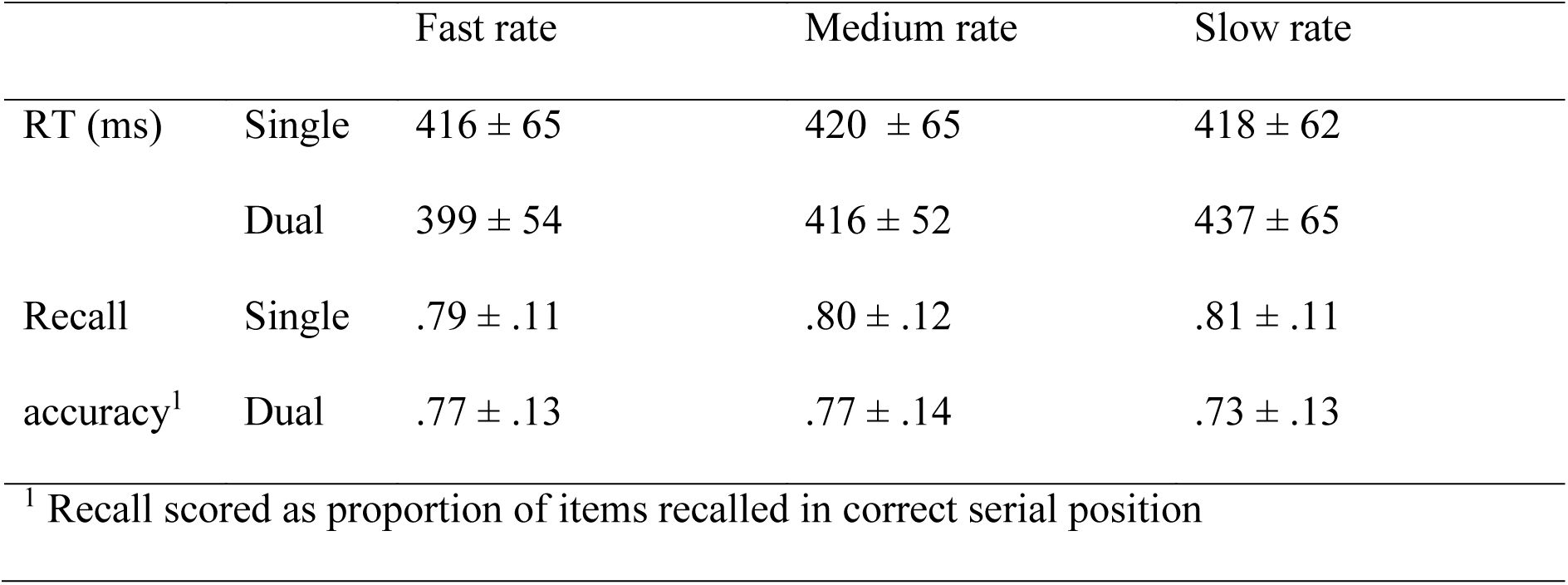
Mean ± SDs for performance in concurrent CRT task and running span task, for each load and rate condition

### Recall data

Recall accuracy in single and dual task was compared across the three rate conditions using a 2×3 repeated measures ANOVA. Attentional load had a significant effect on recall, *F*(1,29) = 26.27, *p* < . 05, 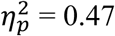, and this interacted with rate, *F*(2,58) = 14.26, *p* < . 05, 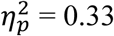. Pairwise Bonferroni comparisons showed the relative impairment in performance in the dual task condition increased with a decrease in task pace, *d* = .45 (fast), = .63 (mid), = 1.06 (slow), *p*s < .05.

To investigate whether there was an impact of rate on the serial position function, a 3×4 repeated measures ANOVA was conducted (Figure 9). For this analysis, recall accuracy was collapsed across the single and dual task condition. There was a strong overall serial position effect, *F*(3,87) = 125.56, *p* < . 05, 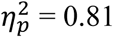, but no interaction with rate, *F*(2,58) = .78, *p* > .05. For each rate condition, a significant 10% improvement in recall accuracy was observed with each consecutive serial position, *p*s < .05.

**Figure 9.**
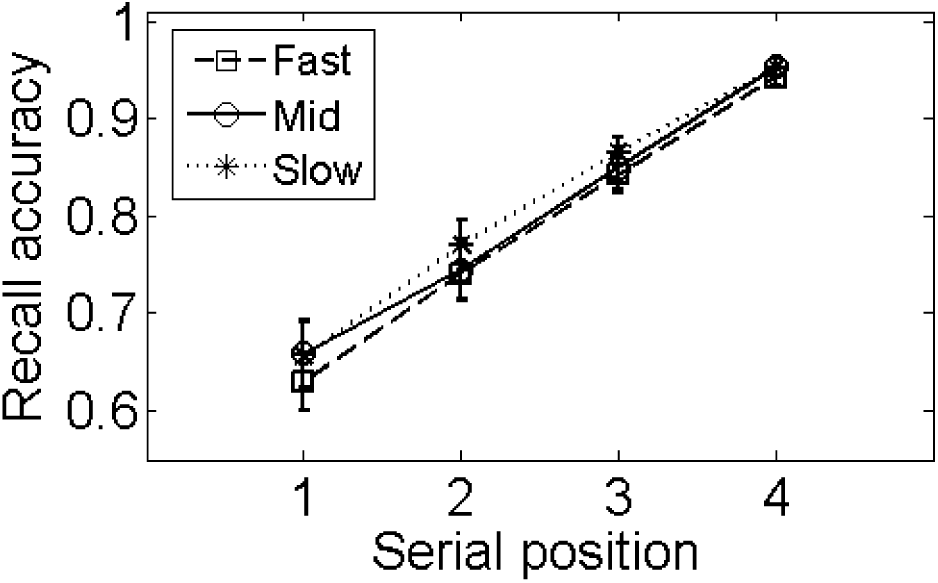
Recall accuracy (proportion of items recalled in correct serial position) as a function of positions in the list. Data depicted for fast (dashed), mid (solid) and slow (dotted) rate, collapsed across the single and dual load within each rate condition.

### Strategy use

Participants reported the fast-paced task as the least demanding condition (N=15), followed by the medium (N=12) and slow paced task (N=3), *χ^2^*(2, N = 30) = 7.8, *p* < .05. The frequencies of strategy use are summarized in Figure 10. Participants reported using fewer strategies to support recall at the fast presentation rate (0.93) compared with the medium (1.13) and the slow rates (1.39). The five active strategies were most frequently employed in the slow rate, followed by the medium and fast rate conditions respectively. The exception to this pattern was passive listening, which was most frequently reported in the fast rate condition. Non-parametric Friedman tests revealed that strategy use differed significantly across rates for all strategies, *p*s < .05, except the use of mental imagery, *p* > .05.

**Figure 10.**
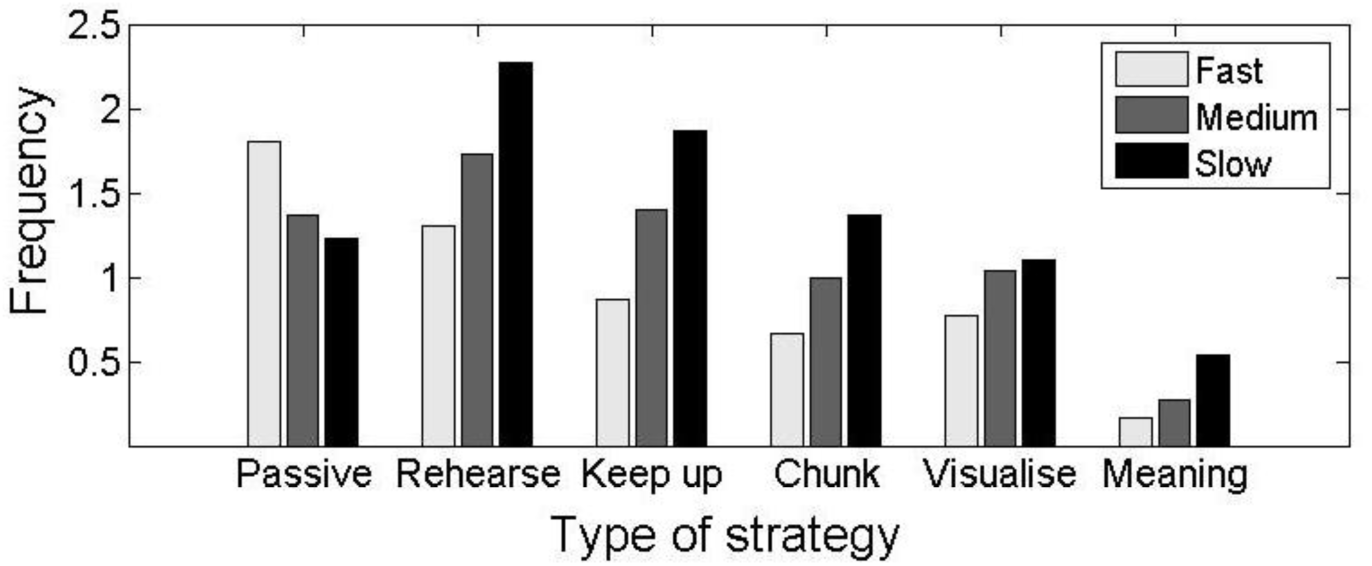
Mean frequency of strategy use self-reported for each rate condition. Participants rated their use of each strategy from 0 (=almost never) to 4 (=almost always).

## Discussion

In line with predictions, the delays in concurrent RTs increased as the presentation rate was slowed. At the fast presentation rate, the RT cost was minimal across all serial positions and did not increase during later positions in the list. At both medium and slow rates, the concurrent RT functions diverged from the fast-paced condition at the third and fourth position respectively and then held constant across most later positions. A localized increase in concurrent RT occurred around 1000ms following the onset of update items (position *n* onward), but only at the slow rate. Participants also reported greater use of active strategies at the slow rate, with passive listening used most commonly at the fastest presentation rate.

Multiple active strategies were reported including rehearsing items, keeping up with the target set, chunking and using mental imagery. Data reported by Hockey (1973) indicated that recall was optimized when participants adopted a passive strategy during fast presentation or an active strategy at slower presentation rates. Bunting et al. (2006) built on these findings by showing that the divergence in strategy use emerges spontaneously in two groups of participants when given the appropriate rate conditions. We extend this further here by demonstrating that the same participants flexibly deployed strategies according to the opportunities afforded by presentation rate.

The peak in resource demands 1000ms after item onset at the slowest presentation rate 1600ms/item in this experiment corresponded that observed at the even slower rate of 2400ms/item in the previous experiment. This invariance indicates a fixed time course for the complex processes involved in updating the target set rather than one that is modulated by the experimental context. Given what looks to be an invariant time lag of one second, it is no surprise that performance suffers when participants attempt to adopt an active updating strategy at faster rates (Hockey, 1973).

Contrary to previous observations, our recall data did not show an effect of rate on serial position functions. Bunting et al. (2006), for example, reported a shallower serial position curve in the slow task compared with the fast task (see too Hockey, 1973). In their investigation, the rate effect was only found at positions four to six from the end of the list, suggesting that the active and passive strategies were equally effectively for the last three positions in the target set. The set size employed here was relatively small because our primary interest was in the processes involved in accurate updating and not recall accuracy. This may be the reason why it was insensitive to subtle changes across intermediate serial positions. A recall advantage in the slow rate condition for the early target items may become evident in lengthier sequences.

## General Discussion

Two experiments tracked the demand on cognitive resources imposed by a verbal running span task requiring the serial recall of the four most recently presented items. When items were presented at a slow pace, running span placed a substantial but specific demand on cognitive resources (Experiment 1 & 2). The cognitive demand in running span emerged around position *n* in the list establishing a heightened baseline that was maintained for the rest of the list. The data also demonstrated localised peaks in demand that occurred around 1000ms following the onset of these later items. This demand profile was sensitive to the presentation rate and disappeared when the rate was faster than one item per second (Experiment 2). We propose that the heightened demand reflects a time and resource-consuming active updating process (Bunting et al., 2006; Cowan et al., 2005; Hockey, 1973; Weems et al., 2009). This cannot be applied at faster rates, thus forcing participants to rely on passive listening instead. The strategy data were consistent with this interpretation.

As yet there is no detailed cognitive model of the processes involved in active updating in running span. One possibility derived from the removal account of updating (Kessler & Oberauer, 2014; Lewis-Peacock, Kessler & Oberauer, 2018; Oberauer, 2018) is that all items in the target set could be actively unbound from their current positions successively. The first item would remain unbound (and thus removed) while the remaining relevant items would then be rebound to new positions allowing the new item to be incorporated into the target set. An alternative account applied to the primacy model of serial recall (Page & Norris, 1998) is that the first item in the target set could get selectively suppressed at each updating episode. This would reset the activation gradient such that the second item would now receive highest activation but maintain the ordered representation of the items. The present data do not distinguish the mechanism but they do establish temporal characteristics to inform these accounts.

Recency but not primacy effects were present in all running span conditions, a finding in line with previous studies (e.g. Bunting et al., 2006; Elosúa & Ruiz, 2008; Hockey, 1973; Morris & Jones, 1990; Palladino & Jarrold, 2008). Ruiz and colleagues have argued that the absence of a primacy effect in running span constitutes evidence against active updating (Elosúa & Ruiz, 2008; Ruiz & Elosúa, 2013; Ruiz, Elosúa, & Lechuga, 2005; see also, Palladino & Jarrold, 2008). They argued that if updating occurred successfully, positional errors in running span should mirror those in serial recall. In other words, a higher proportion of errors was expected in medial serial positions in running span, generating standard primacy and recency effects. On this basis, it was concluded that the recency effect in running span task reflected passive listening (see also Broadway & Engle, 2010). These proposals are inconsistent with our findings. The concurrent RT data presented here clearly demonstrate that resource consumption in running span is sensitive to the rate of presentation, indicating a different set of cognitive processes at slow and fast rates. Our investigation highlights the need for alternative measures; interpretations of end-of-list recall data may be unable to capture cognitive processing during list presentation.

Hockey and Hamilton (1977) put forward an alternative explanation for the lack of primacy in running span. Their suggestion was that that primacy in standard serial recall originates in a perceptual advantage associated with the start of the target list. In running span, items preceding the current target set would eliminate this advantage. Their hypothesis is supported by the emergence of a primacy effect in running span when the first target item in an ordered sequence is either the first item presented of the list (Hockey & Hamilton, 1977) or is signaled by a perceptually distinctive cue (Ruiz & Elosúa, 2013). These are conditions under which an updating operation may not be necessary to keep track of the relevant target items, as indeed in our modified span task in Experiment 1. Applying these conditions may diminish or even possibly erase the updating-related demand observed in the present study. Indeed, the conventional absence of primacy in running span may be indicative of the presence, rather than the absence, of an update process.

While our data clearly chart the time course of updating in running span under concurrent task conditions, a fuller understanding of the complexity of updating in working memory requires substantial further investigation. One key issue is whether the time course of updating seen in the present experiments is itself influenced by the requirement to perform a concurrent task or is genuinely fixed. We also acknowledge that there are other important forms that updating in working memory can take other than the particular form of serial updating involved in running span. New work is required to establish whether the time course of updating changes when task demands change, for example with the serial recognition requirements of n-back (e.g. Jaeggi, Buschkuehl, Perrig & Meier, 2010; Kane, Conway, Miura, & Colflesh, 2007) or updating based on non-serial item characteristics such as semantic category or spatial location (Kessler & Oberauer, 2014). We also know little about the apparently heterogenous set of active strategies reported by participants in the slow-paced running span conditions of Experiment 2 and whether their temporal characteristics can be distinguished. What we can conclude with reasonable confidence is that the 1000ms peak on resource demand is robust in this verbal serial updating paradigm.

